# Circadian Control of Pulmonary Endothelial Signaling occurs via the NADPH oxidase 2-NLRP3 pathway

**DOI:** 10.1101/2022.06.05.493624

**Authors:** Shaon Sengupta, Yool Lee, Jian Qin Tao, Amita Sehgal, Shampa Chatterjee

## Abstract

Circadian rhythms are endogenous oscillations that occur with a 24-hr periodicity. These rhythms are ubiquitous and thus, vascular endothelial cells that line the vascular bed are also subjected to circadian regulation. While the circadian control of vascular function has been demonstrated in the context of various pathologies, the relevance and functional implication of clock control over pulmonary vasculature has never been investigated. As the pulmonary endothelium is a crucial site for the host’s inflammatory response to a lung specific pathogen, we investigated the role of the circadian clock in mediation the response of the pulmonary endothelium to inflammation. We hypothesized that the pulmonary endothelium is under circadian control and that the clock serves to curb inflammatory signaling.

**Methods:** Circadian rhythms were monitored in pulmonary artery segments and endothelial cells isolated from mPer2luciferase transgenic mice in the presence of an inflammatory stimuli (LPS). Reactive oxygen species (ROS) production in LPS treated cells was measured by fluorescence microscopy using the cell permeant dye CellROX Green. NLRP3 inflammasome was monitored post-mortem (0-72 h post LPS instillation) by measuring the expression of the NLRP3 subunit in wild type and *Bmal1^−/−^* and *Cry1/2*^*−/−*^ mice. Inflammation was quantified in these mice by measuring PMN adherence and intercellular adhesion molecule (ICAM-1).

**Results:** We observed that the circadian rhythm of the pulmonary vasculature was altered LPS. LPS also led to ROS production in these cells; ROS increased 3 h post LPS treatment, peaked by 36 h and returned to baseline values by 72 h. ROS were inhibited by pretreating the cells with the NADPH oxidase 2 (NOX2) inhibitor dipheneylene iodonium (DPI). Addition of DPI, prior to LPS pretreatment also restored the circadian rhythmicity of the pulmonary endothelium. The increase in NLRP3 along the vessel wall (post LPS treatment) was resolved by 72 h in lungs of wild type mice but not in *Bmal1^−/−^* and Cry1^−/−^Cry2^−/−^ lungs. Inflammation (ICAM-1 and PMN) was also resolved in wild type but not in mice wherein the circadian clock had been disrupted genetically.

**Conclusion:** Our data indicate that pro-inflammatory stimuli reprogram circadian rhythms in the pulmonary endothelium via ROS via the NOX2-NLRP3 pathway. Disruption of the clock mediates a sustained increase in ROS via this Nox2-NLRP3 pathway in endothelial cells, thus offering a novel mechanism for mitigating the effects of clock disruption.

## Introduction

Circadian rhythms refer to endogenous oscillations that occur with a 24-hr periodicity which and are manifest in almost all cells [1]. Studies on the systemic vascular endothelium (a network that integrates biochemical and biophysical signals via the transport of blood, nutrients, inflammatory and pathogenic stimuli) have established that endothelial cell signaling is subject to circadian regulation [2–4]. Indeed, the circadian clock has been characterized in the vasculature of systemic organs and endothelial signaling along with inflammation has been found to show circadian regulation [5, 6]. However, the circadian regulation of lung endothelium has not been studied. As the lung forms the first line of defense to air borne pathogens [7] and the pulmonary endothelium by virtue of its location is pivotal in onset and progression of the inflammatory response [8, 9], the circadian influence on pulmonary endothelium needs to be investigated. Our earlier work reported that the lung endothelium responds to endotoxin exposure (LPS administration) by assembly of the endothelial NADPH oxidase 2 (NOX2) leading to the production of reactive oxygen species (ROS) [10]. The ROS thus produced activated inflammation signaling [11].

We aimed to investigate the biological significance of the endogenous clock in the inflamed pulmonary endothelium. We hypothesized that lung endothelial signaling is subject to circadian control which is regulated by NOX2 induced ROS. We investigated (1) The effect of inflammation on the circadian rhythms of the pulmonary endothelium and the effect of NOX2 blockade on the same and (2) Whether the heightened inflammation seen in the pulmonary vasculature of clock disrupted mice is mediated by the NLRP3 inflammasome pathway. Our findings indicate that the circadian clock is affected by inflammatory signaling, and its presence serves to mitigate pulmonary inflammatory damage.

## Materials and Methods

### Materials

LPS (O111:B4) and apocynin were from Sigma-Aldrich (St Louis, MO). Antibodies against ICAM-and NLRP3 inflammasomes were purchased from BD Biosciences. Dynabeads® were from Dynal (Oslo, Norway) pre-labeled with sheep anti-rat IgG. Animal use was reviewed and approved by the University of Pennsylvania Institutional Animal Care and Use Committee.

### Mice

mPer2-luciferase on C57Bl6 background were a kind gift of J Takahashi, UTSW, Texas. *Bmal^1−/−^* mice were on a C57Bl6 background (???) [12] CRY 1/2 double knockout mice were a gift from Katja Lamia originally from Aziz Sancar [13]. Age and background matched wild type littermates were used as control. All animal studies were approved by the University of Pennsylvania Institutional Animal Care and Use Committee and met the stipulations of the Guide for the care and Use of Laboratory animals.

### Methods

#### Lung explants and isolation of pulmonary artery from mPer2-luciferase mice

Mice were anesthetized, and a tracheostomy was performed. Lungs were ventilated and perfused in Krebs-Ringer buffer to remove blood. Perfusion-cleared lungs were excised from the chest cavity and chopped into cubes of 0.5 mm. In separate experiments, a section of the pulmonary artery was removed from the excised lungs. For this, the lungs were dissected with micro-scissors (to remove superficial tissue) using a stereomicroscope (Leica Microsystems, Buffalo Grove, IL) and the main pulmonary artery removed. The adventitia was separate from the isolated arteries and extreme care was taken to avoid any damage or stretching.

#### Isolation of pulmonary microvascular endothelial cells (PMVEC)

Endothelial cells were isolated from the lungs of these mice as described by us previously ([14, 15]. Briefly, lungs from mPer2-Luc mice are removed and the tissue trimmed at the periphery. The trimmings were treated with collagenase (3 mg/ml) and the digest pelleted and incubated with monoclonal anti-PECAM antibody. This was followed by addition of magnetic beads (Dynabeads®, Dynal, Oslo, Norway) pre-labeled with sheep anti-rat IgG. The beads were separated on a magnetic rack, washed and plated on culture plates. Isolated endothelial cell islands were obtained in 1-2 weeks. A second round of immunoselection is carried out by flow sorting PECAM positive cells by labeling the cells with anti-PECAM-FITC. The endothelial phenotype of the preparation was confirmed by evaluating cellular uptake of the endothelial specific marker such as DiI-acetylated low-density lipoprotein (DiIAcLDL) and immunostaining for platelet endothelial cell adhesion molecule (PECAM-1), lung tissue was trimmed at the periphery, digested and PECAM positive cells isolated by immunoselection using magnetic beads [15].

#### Circadian Oscillations

Pulmonary endothelial cells of mPer2Luc were treated with 50 µg LPS and seeded in 35 mm dishes (at 80% confluence). Lung explants and pulmonary arteries were excised from perfused lungs for each mouse. Arteries were carefully cleaned from surrounding connective tissue. 4-5 mm pieces of explant or segments of 5 mm cut from each artery, were placed in tissue culture dishes and treated with LPS. Real time oscillations of the circadian gene Per2 promoter activity were assessed by recording the bioluminescence (Lumicycle32, ActiMetrics) as described previously [16]. Data was analyzed using LumiCycle software (Actimetrics).

#### Endotoxin Instillation

Mice were anesthetized by intraperitoneal (IP) injection of ketamine (100 µg/g body wt), xylazine (4 µg/g) and acepromazine (1 µg/g). LPS (5 mg/kg per mouse; serotype O111:B4; Sigma, St. Louis, MO) dissolved in 50 μl sterile 0.9% NaCl was instilled intratracheally (IT) via a canule, followed by 0.15 ml of air. After IT treatment, the mice were kept in an upright position for 10 min to allow the fluid to spread throughout the lungs. Mice were euthanized at 6-72 h after instillation. Control mice (0 time) received PBS.

#### Reactive oxygen species (ROS)

was monitored in LPS and vehicle treated pulmonary microvascular endothelial cells (PMVEC) by labeling cells with CellROX® as described earlier [17, 18]. Briefly, PMVEC were incubated for 30 min with 5 μM Cell ROX and imaged by epifluorescence microscopy using a Nikon TMD epifluorescence microscope, equipped a Hamamatsu ORCA-100 digital camera, and Metamorph imaging software (Universal Imaging, West Chester, PA, USA). Images were acquired at λex =488 nm using both 4X and 20X lenses. All images were acquired with the same exposure and acquisition settings as reported previously [17, 19].

#### Inflammation

The expression of the NLRP3 subunit of the inflammasome expression of ICAM-1 and abundance of polynuclear neutrophils (PMN) were monitored at various time periods post-LPS instillation by immunostaining. Mice were sacrificed, lungs perfused, excised, and fixed in 4% paraformaldehyde. This was followed by dehydration by sequential sucrose treatment (10, 20 and 30% sucrose). The tissue was embedded in OCT blocks and longitudinal sections were immune-stained for ICAM-1, NLRP3 and Ly6G by using monoclonal anti-ICAM, NLRP3 antibody, and polyclonal NIMP respectively. Secondary antibodies used were coupled to Alexa 488. Images were acquired on a Nikon Fluorescence (Nikon Diaphot TMD, Melville, NY). Appropriate controls were non-immune IgG and no-antibody treated samples. Images were acquired at excitation of 488 nm at the same settings and the fluorescence quantified over 3-4 fields using Metamorph Software (Molecular Devices, Downington PA). For quantitation of the vascular NLRP3 and ICAM-1, the fluorescent signal along the lung vascular walls is measured using outlining tools of the Software. Data were expressed as arbitrary fluorescence units. Data were expressed as mean ± SD and one-tailed paired t-tests were used to determine statistical significance. Image acquisition and data analysis were carried out in a blinded fashion.

## Results

### Inflammatory Stimulus (LPS) alters the circadian rhythm in lung tissues, pulmonary arteries and pulmonary endothelial cells

We evaluated the effect of lung inflammation on circadian rhythm by exposing explants from freshly isolated lungs (of mPer2-luciferase mice) to LPS. Prior to this, the explants were synchronized with dexamethasome (Dex). Addition of LPS to lung explants altered the circadian rhythm as determined by mPer2luciferase activity (**Figure 1A**). Analysis of the rhythm showed that LPS significantly increased the circadian amplitude **(Figure 1B**) although the period was not affected. Next, to evaluate the effect of inflammation in pulmonary vessels on circadian rhythm, pulmonary arteries were dissected out of these mice and exposed to LPS. Once clocks were synchronized, arteries were found to show a significant increase in amplitude and decrease in period (**Figures 2A, B**).

**Figure 1.**
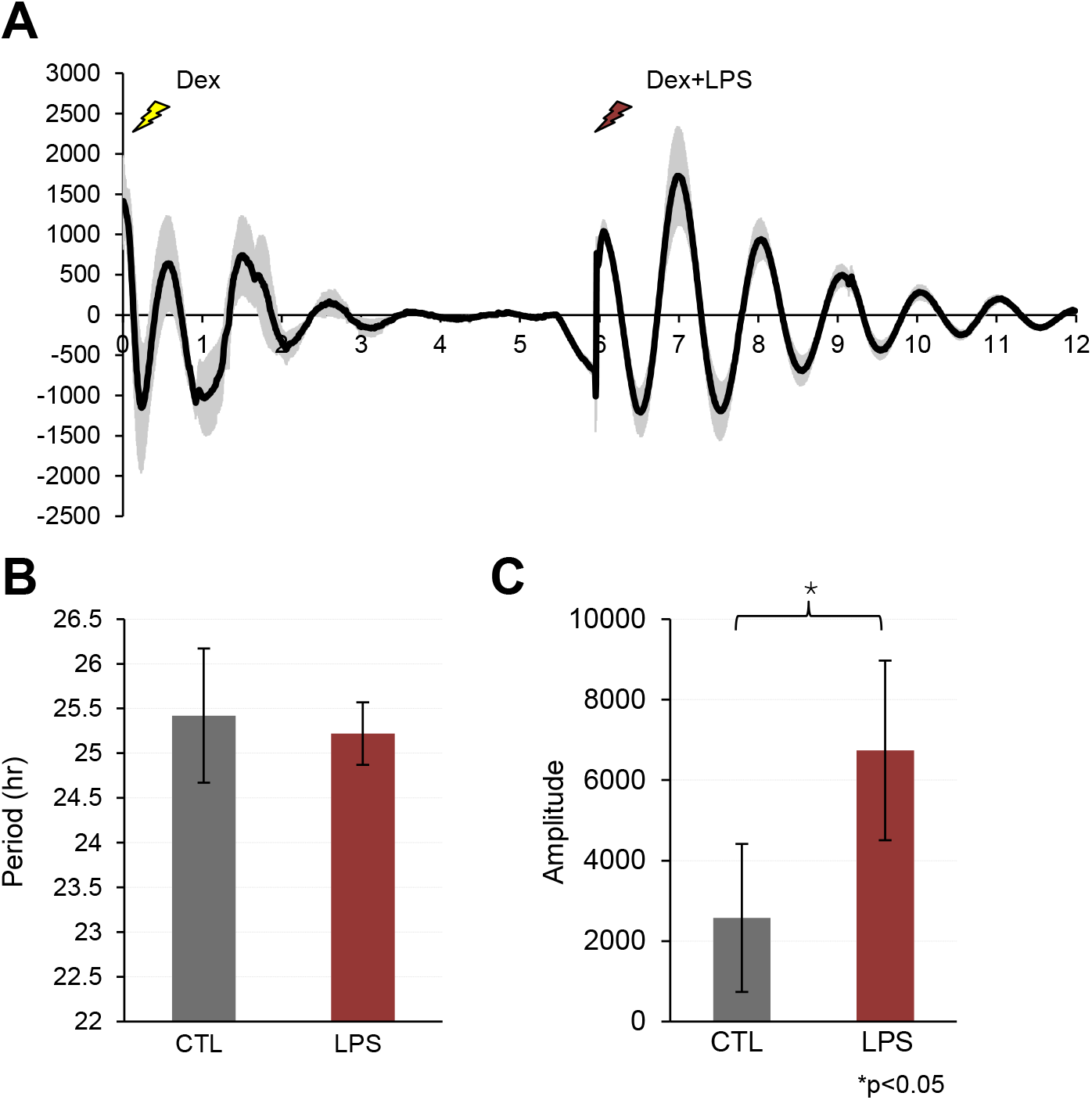
LPS alters the circadian rhythm in synchronized lung explants. A. Representative graphs of bioluminescence data from PER2::LUC tissue explants monitored in culture for 12 days. B. Circadian Period with and without LPS treatment C. LPS increases the amplitude of rhythms.

**Figure 2.**
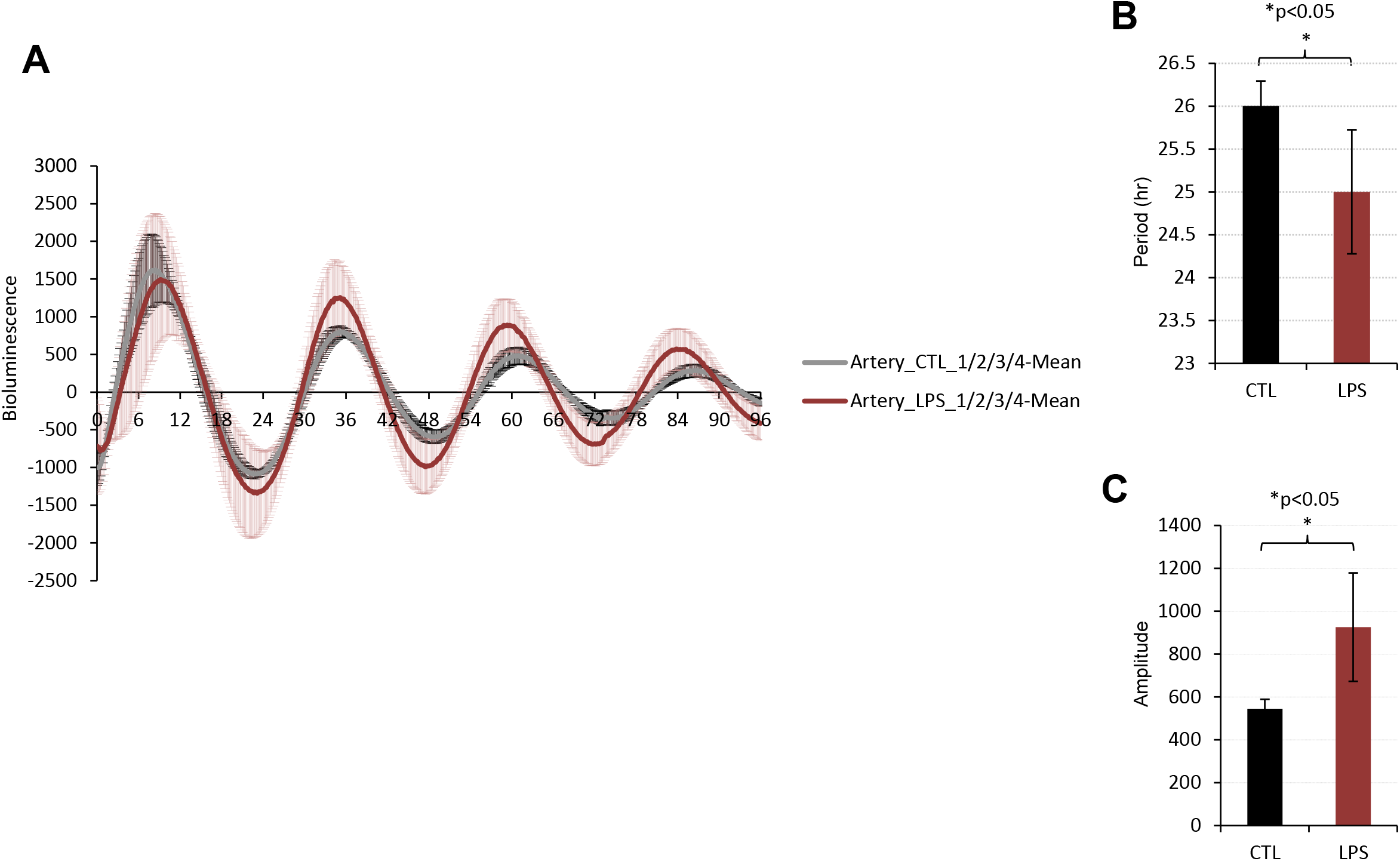
LPS alters the circadian rhythm in dissected pulmonary artery that are synchronized with Dex. A. Representative graphs of bioluminescence data from PER2::LUC pulmonary artery with and without LPS. B. LPS decreases the circadian period C. LPS increases the amplitude of rhythms.

### Blocking ROS induced by inflammation restores circadian rhythm in pulmonary endothelial cells

As the endothelium is a major player in pulmonary vascular function, we investigated the pulmonary endothelial cells for circadian rhythm. Endothelial cells isolated from mPerLuc mice were expanded into large primary cultures. Thus, pulmonary endothelium shows robust circadian rhythms that are reprogrammed in the presence of LPS (**Figure 3**). We have reported earlier that LPS exposure led to the production of ROS in pulmonary endothelial cells [10, 20] and that blocking NOX2 assembly reduced ROS production. Thus, we blocked NOX2 activation by the use of inhibitor apocynin and monitored circadian rhythms in PMVECs treated with LPS. Pre-treating PMVEC with apocynin 1 h prior to LPS treatment blocked ROS production (**Figure 4A**,**B**) and restored the amplitudes of the circadian cycle to control (naïve) values (**Figure 3**), although the circadian period was not restored and showed a further decrease.

**Figure 3.**
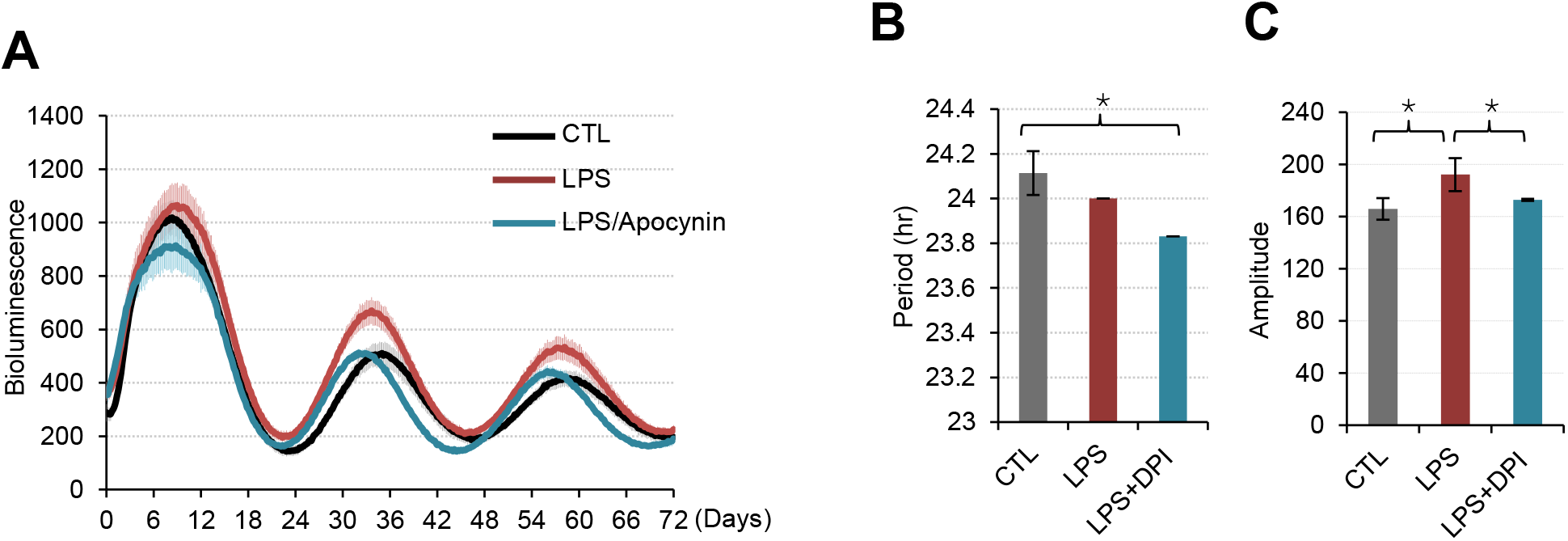
Circadian rhythm in pulmonary microvascular endothelial cells (PMVEC). PMVEC were isolated from lungs of Per2Luc mice. Cells were expanded and endothelial lineage characterized to obtain primary cultures. A. Naïve cells and LPS treated cells that are synchronized show a circadian oscillation as monitored from the luminescence of the reporter. However, LPS induces alteration in circadian rhythmicity of synchronized PMVEC. A phase shift and an increased amplitude is induced with LPS treatment of PMVEC. Treatment with NADPH oxidase 2 inhibitor apocynin resulted in restoration of the amplitude. B. LPS decreases the period and C. LPS treatment increases the amplitude of rhythms which is restored with DPI treatment. *p<0.05

**Figure 4A.**
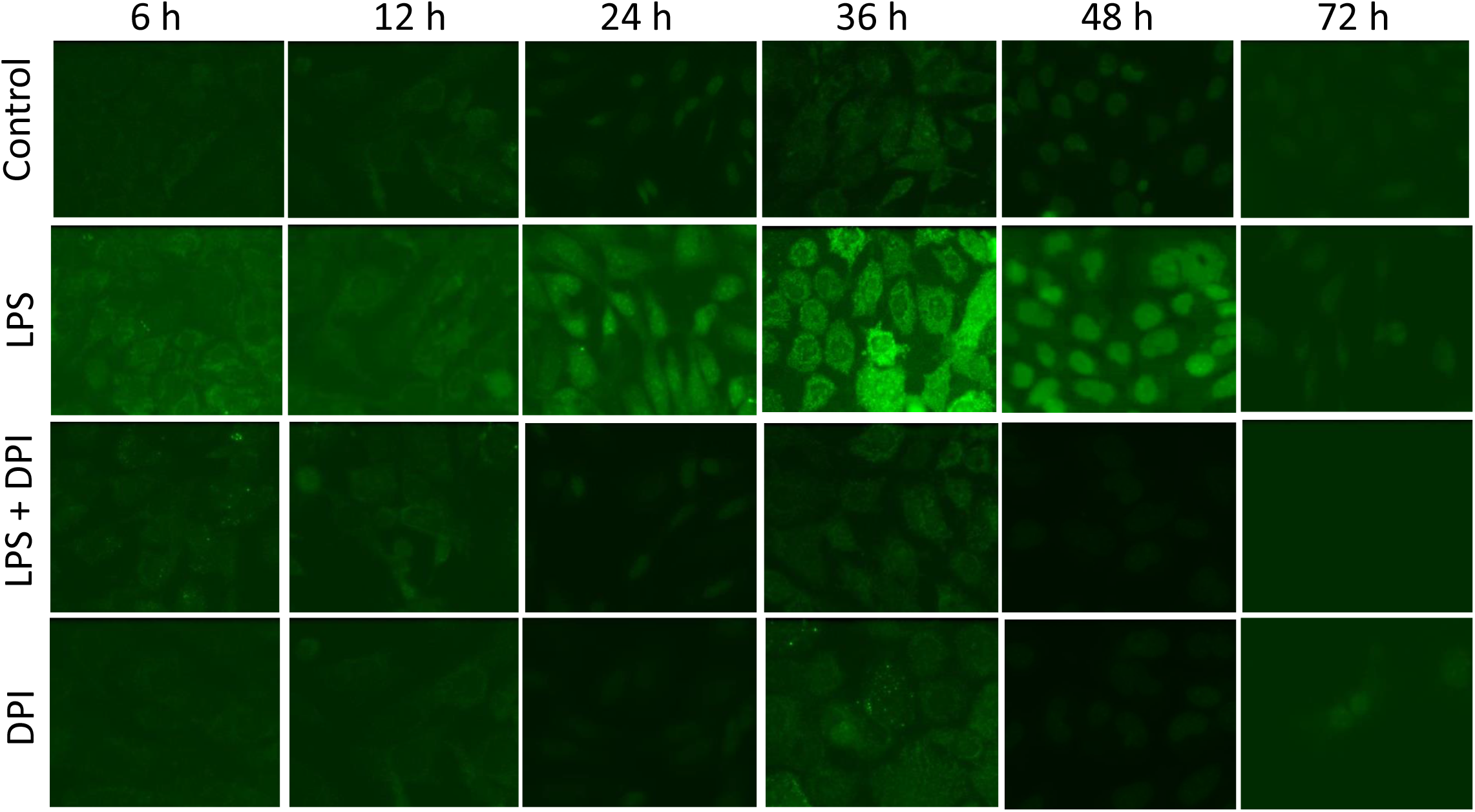
LPS treatment causes ROS production in Per2Luciferase pulmonary endothelial cells. A. Cells were labeled with ROS sensitive dye Cell ROX (5 µM) for 20 min and imaged at λ (ex) by confocal microscopy.

**Figure 4B.**
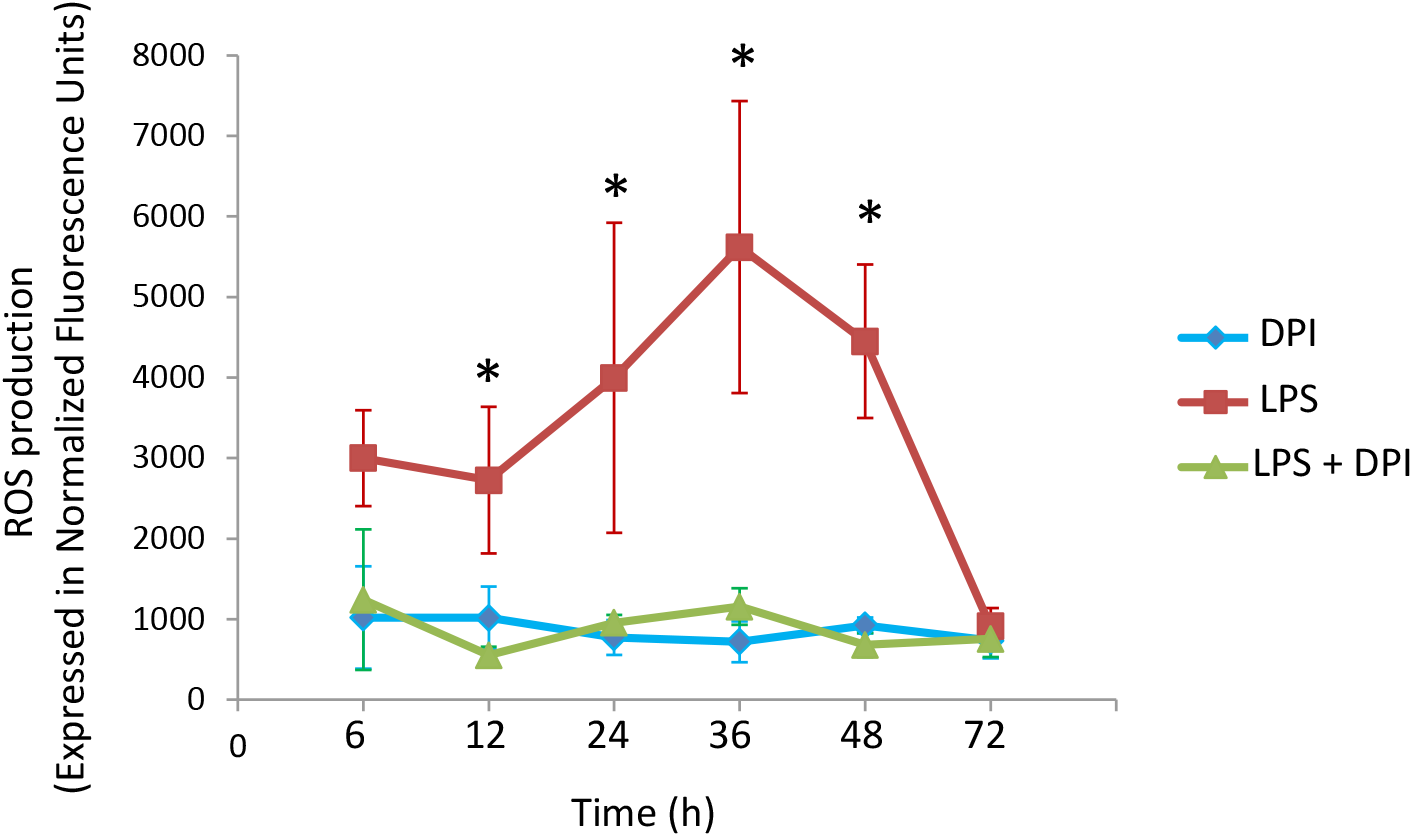
The fluorescence intensity of the images were integrated using MetaMorph Imaging Software. After a gradual increase until 36 h, ROS levels decrease to baseline values. ROS production post-LPS is significantly high as compared to untreated controls and DPI pre-treated cells. *p<0.01 as compared to LPS+ DPI and DPI alone values.

### Circadian Regulation of the endothelial NLRP3 inflammasome

To identify the signals via which ROS regulates circadian rhythm, we evaluated the circadian influence on the endothelial NLRP3 inflammasome. This inflammasome is a master regulator of inflammation. NLRP3 inflammasome in the pulmonary endothelium can be assessed by monitoring the expression of the NLRP3 (the NLRP3 subunit of the inflammasome) along the vessel walls. NLRP3 expression was quantitated along the vessel wall by outlining the fluorescent signal along the vessel wall as we have reported earlier [21].

In mice, we observed the expression of NLRP3 was very low in untreated naive lungs but increases dramatically post-LPS treatment peaking at 36 h and decreasing thereafter to baseline levels (**Figure 5**). Next, we wanted to test if this NLRP3-NOX2 driven ROS signaling is preserved in the absence of an intact clock. For this, we used two models of clock disruption involving the positive and negative limbs of the core clock i.e., *Bmal1^−/−^* and *Cry1,2*^*−/−*^ we found that that this regulation was not observed in the lungs obtained from *Bmal1^−/−^* and *CRY1/2*^*−/−*^ (**Figure 6 A, B**) mice. In these lungs, the NLRP3 expression was high even in the untreated state and did not change significantly upon exposure to LPS (**Figure 6**). Overall, this suggests that disruption of the circadian clock results in a heightened inflammation in the pulmonary endothelium via the NLRP3-NOX2 driven ROS pathways.

**Figure 5A.**
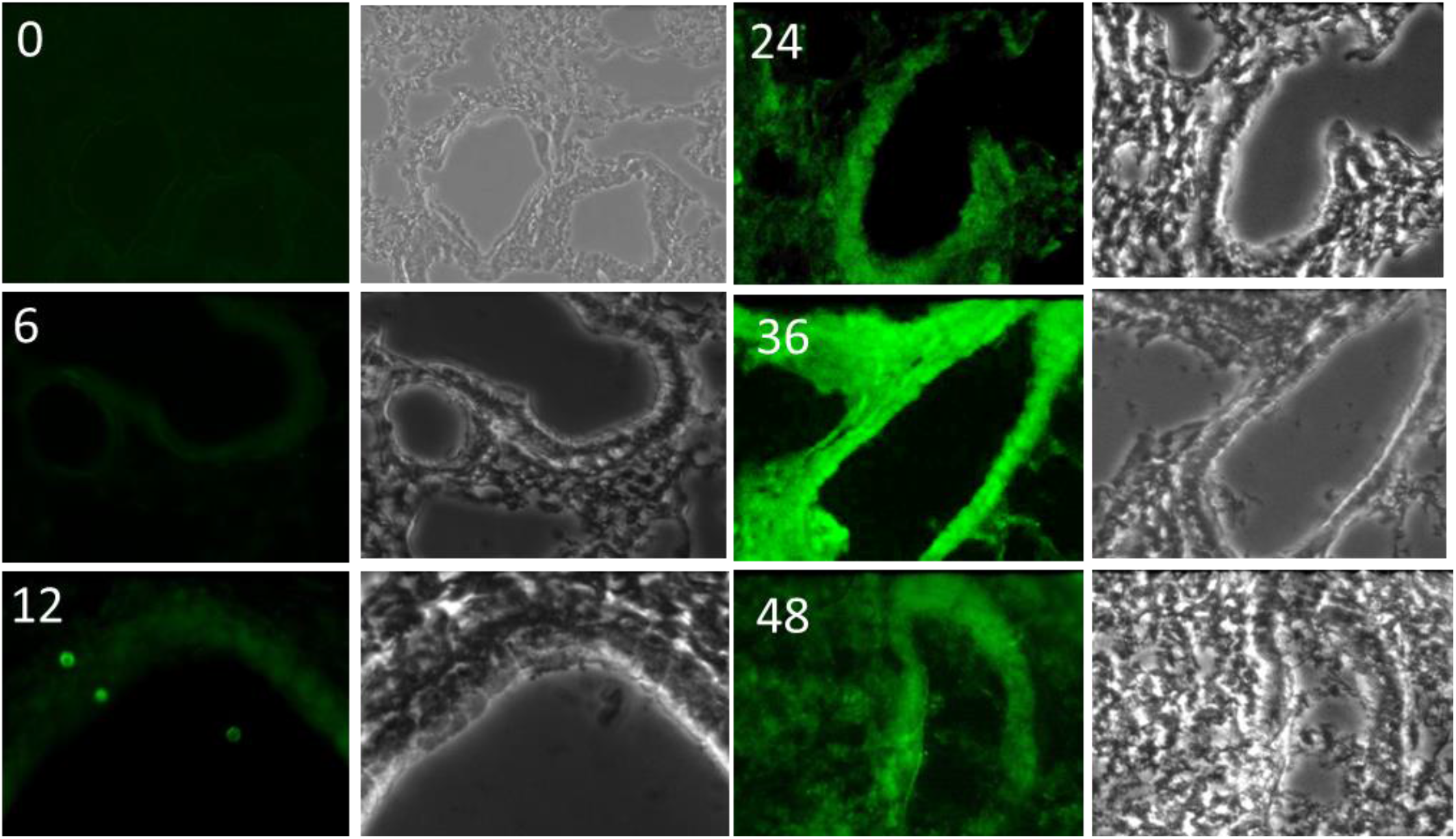
Inflammasome (NLRP3 subunit) expression is induced by LPS. A. mPerLuc mice treated with LPS (intratracheal instillation) were sacrificed at 6-72 h and lungs cleared of blood, fixed and sectioned. Sections were immunostained with anti-NLRP3 antibody. Secondary Antibody was tagged to Alexa 488.

**Figure 5 B.**
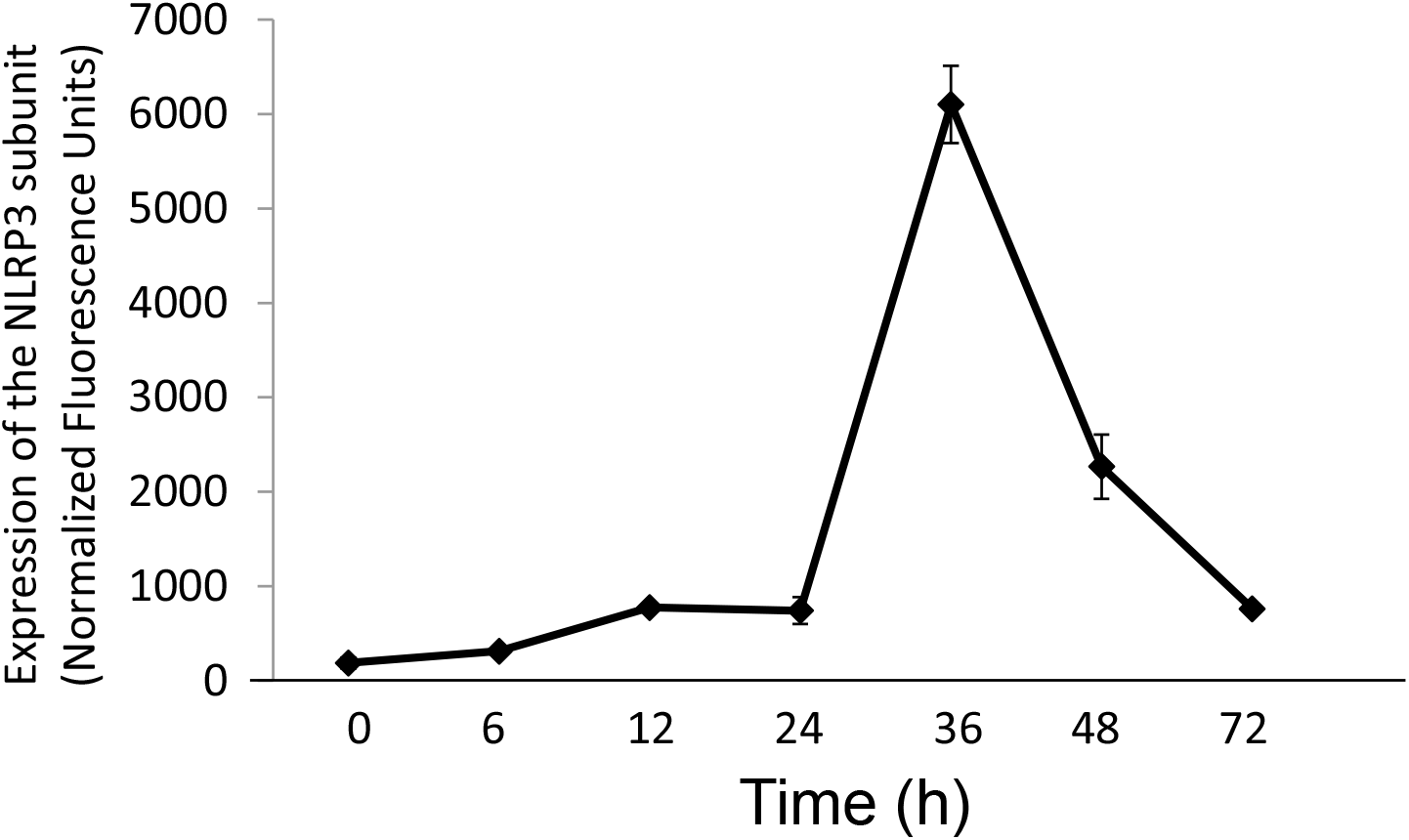
Quantitation of the NLRP3 subunit of the inflammsome. The fluorescence intensity of the images were integrated using MetaMorph Imaging Software.

**Figure 6A.**
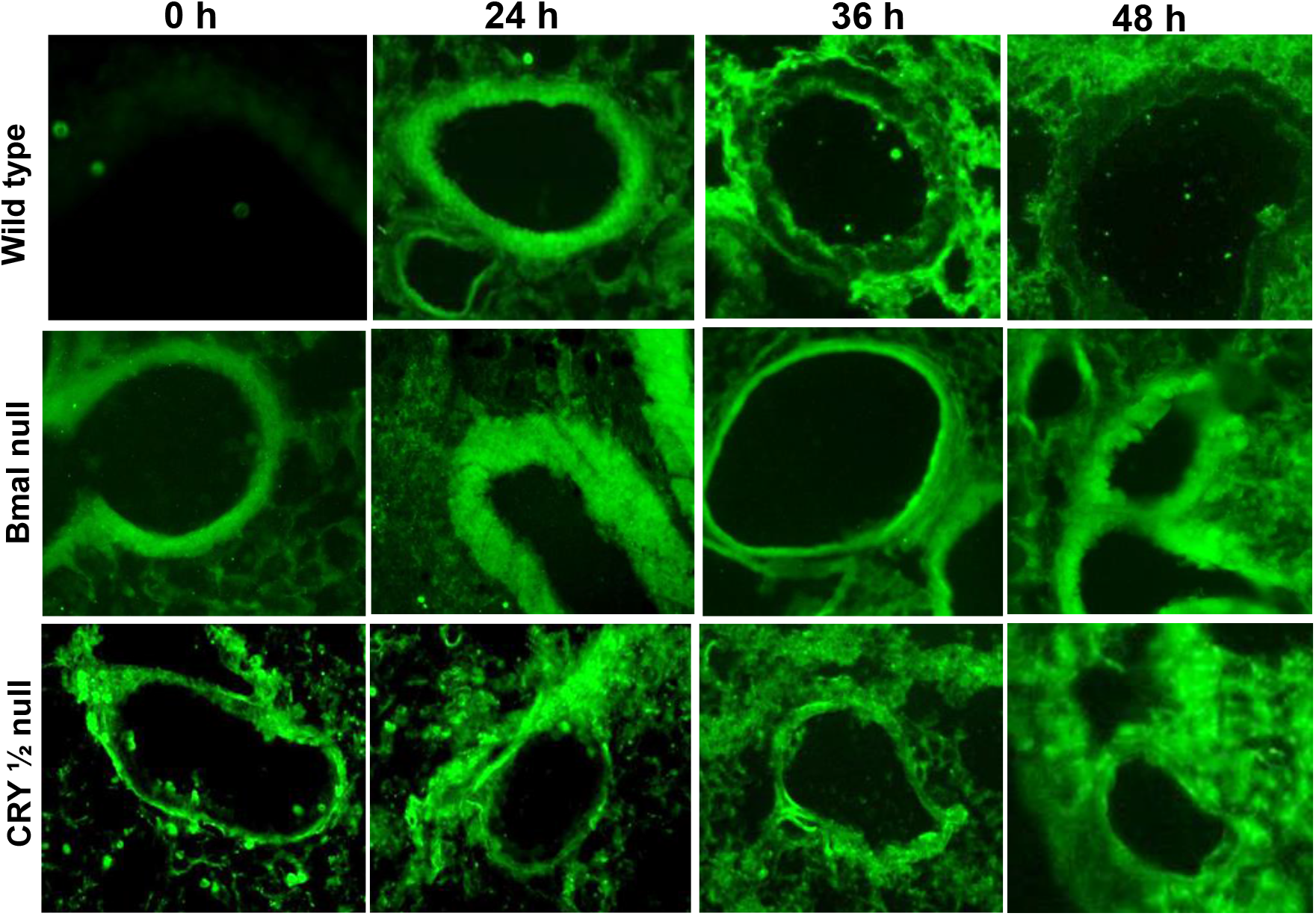
Inflammasome (NLRP3 subunit) expression in WT, Bmal1 null and CRY1/2 null mouse treated with LPS. Mice were sacrificed at 24-48 h and lungs cleared of blood, fixed and sectioned. Sections were immunostained with anti-NLRP3 antibody. Secondary Antibody was tagged to Alexa 488.

**Figure 6B.**
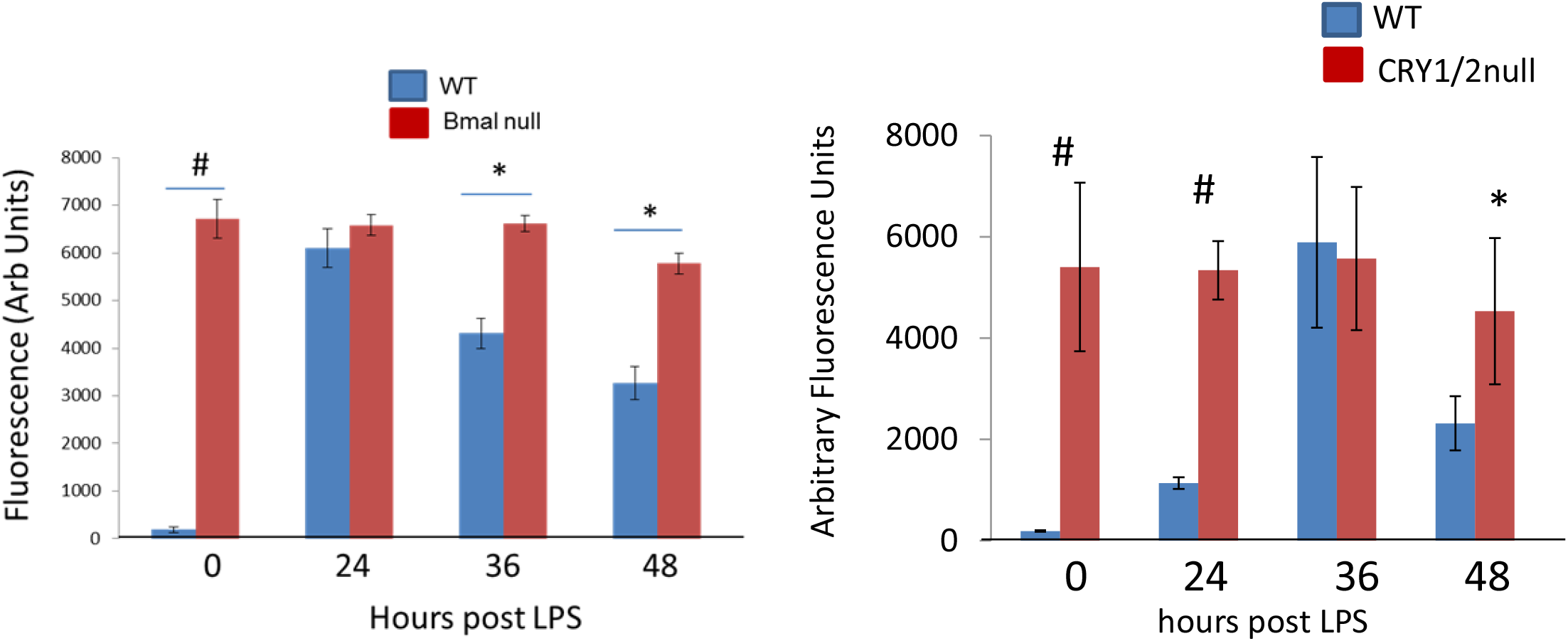
Quantitation of the NLRP3 subunit of the inflammasome. The fluorescence intensity of the images were integrated using MetaMorph Imaging Software. *p<0.05, #p<0.001

### Circadian Regulation of inflammation (ICAM-1 and PMN influx)

Inflammation was monitored by measuring expression of cellular adhesion molecule specifically, intercellular adhesion molecule-1 (ICAM-1) and adherence and extravasation of polymorphonuclear neutrophils (PMN). In wild type mice, LPS instillation led to a gradual increase in ICAM-1 along the lung vessels and a decrease thereafter. Such regulation was not observed in Bmal1^−/−^ and CRY1/2^−/−^ mice where ICAM-1 was highly expressed even in the naïve untreated lungs and did not significantly change with LPS exposure (**Figure 7**). PMN that adhered/extravasated into lung tissue were imaged (**Figure 8A**) and assessed by either a) counting the number of positive signals using Ly6C stained lung sections (**Figure 8B**) or b) total fluorescent signaling emanating from the sections to represent activated PMN (**Figure 8C**). Total adhered and extravasated PMN in lungs were low in naïve untreated lungs but increased with LPS instillation. With Bmal1 and CRY1/2 deletion, PMN was high in naïve lungs and did not increase appreciably with LPS exposure. Thus, ablation of the circadian clock (Bmal1 and CRY1/2) prevented the regulation of inflammation stimulus.

**Figure 7A.**
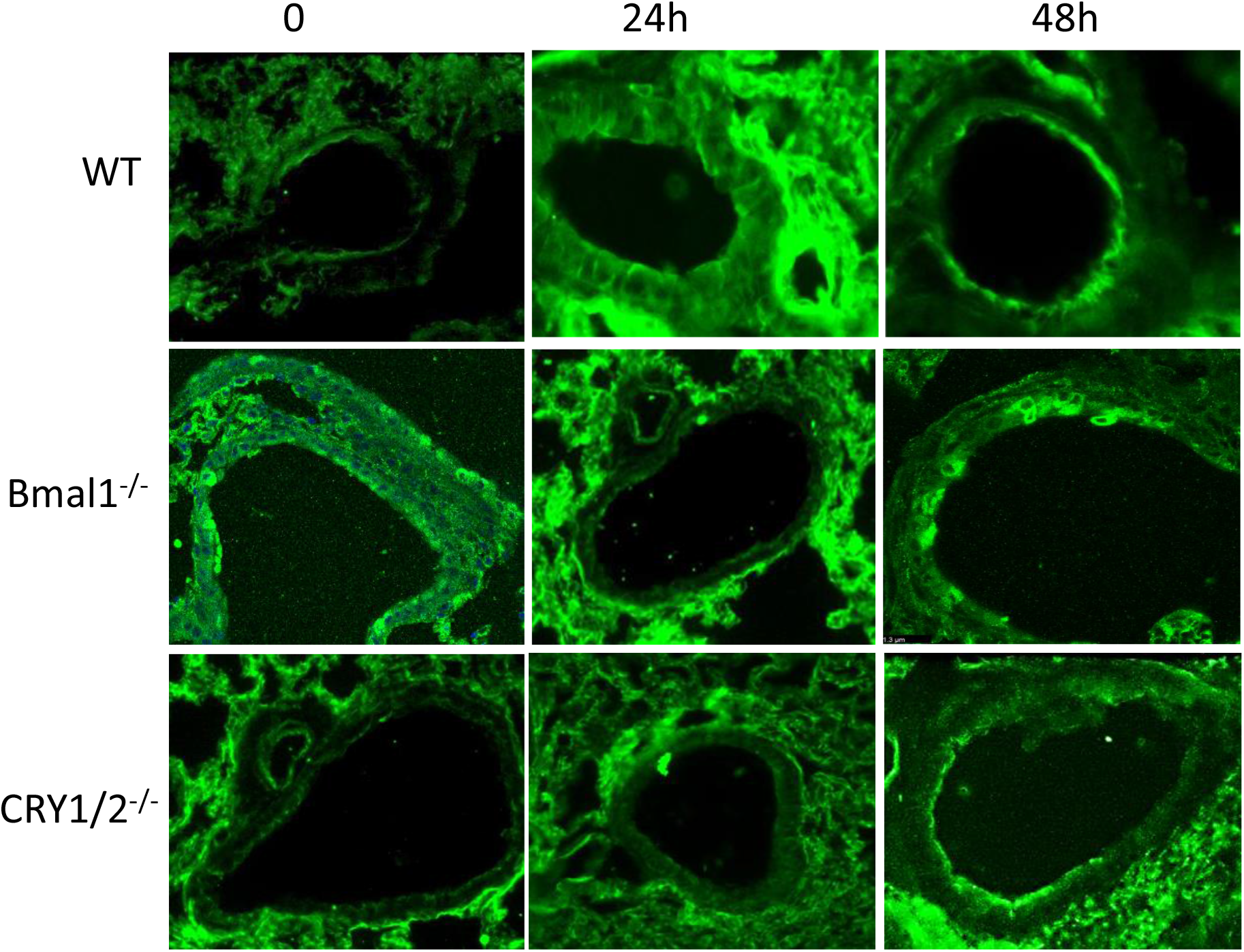
Inflammation in lungs as monitored by expression of intercellular adhesion molecule (ICAM-1) in WT, Bmal null, CRY1/2 null mouse treated with LPS. Mice were sacrificed at 24-48 h and lungs cleared of blood, fixed and sectioned. Sections were immunostained with anti-NLRP3 antibody. Secondary Antibody was tagged to Alexa 488.

**Figure 7B.**
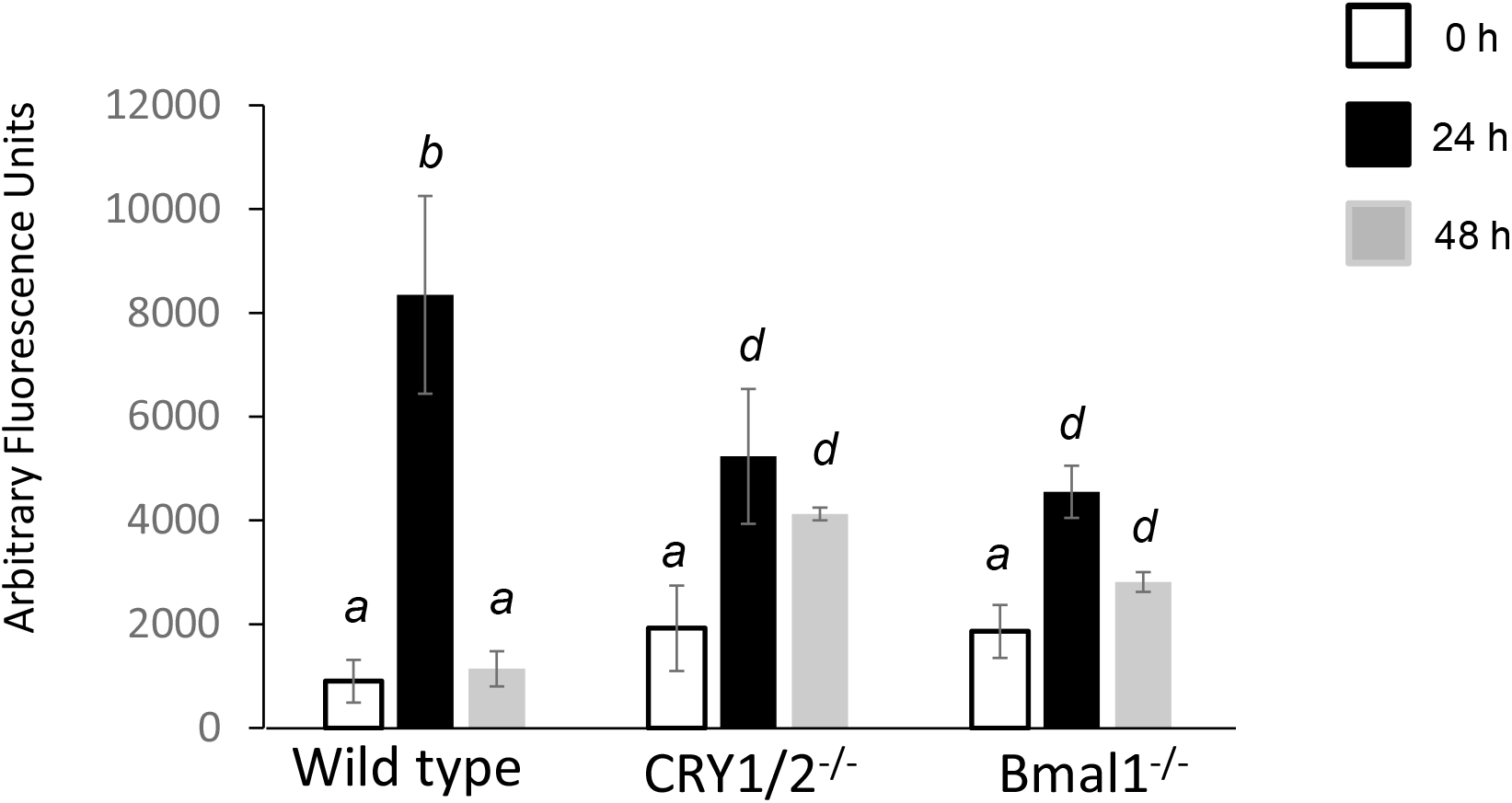
Quantitation of ICAM-1. The fluorescence intensity of the images were integrated using MetaMorph Imaging Software. Fluorescence signal (green signals per field) was quantitated over several sections using Metamorph Software. n=3 WT mice and CRY1/2 null and n=2 Bmal null were used. For each mouse, at least 4-5 sections were immunostained and imaged. 3 fields were imaged per section to obtain an average for each section. All sections was averaged, and data as expressed as means± SD. Different letters between groups denote statistically significant differences (p<0.05) between those two groups. If two groups share the same letter, then they are not statistically significantly different

**Fig 8A.**
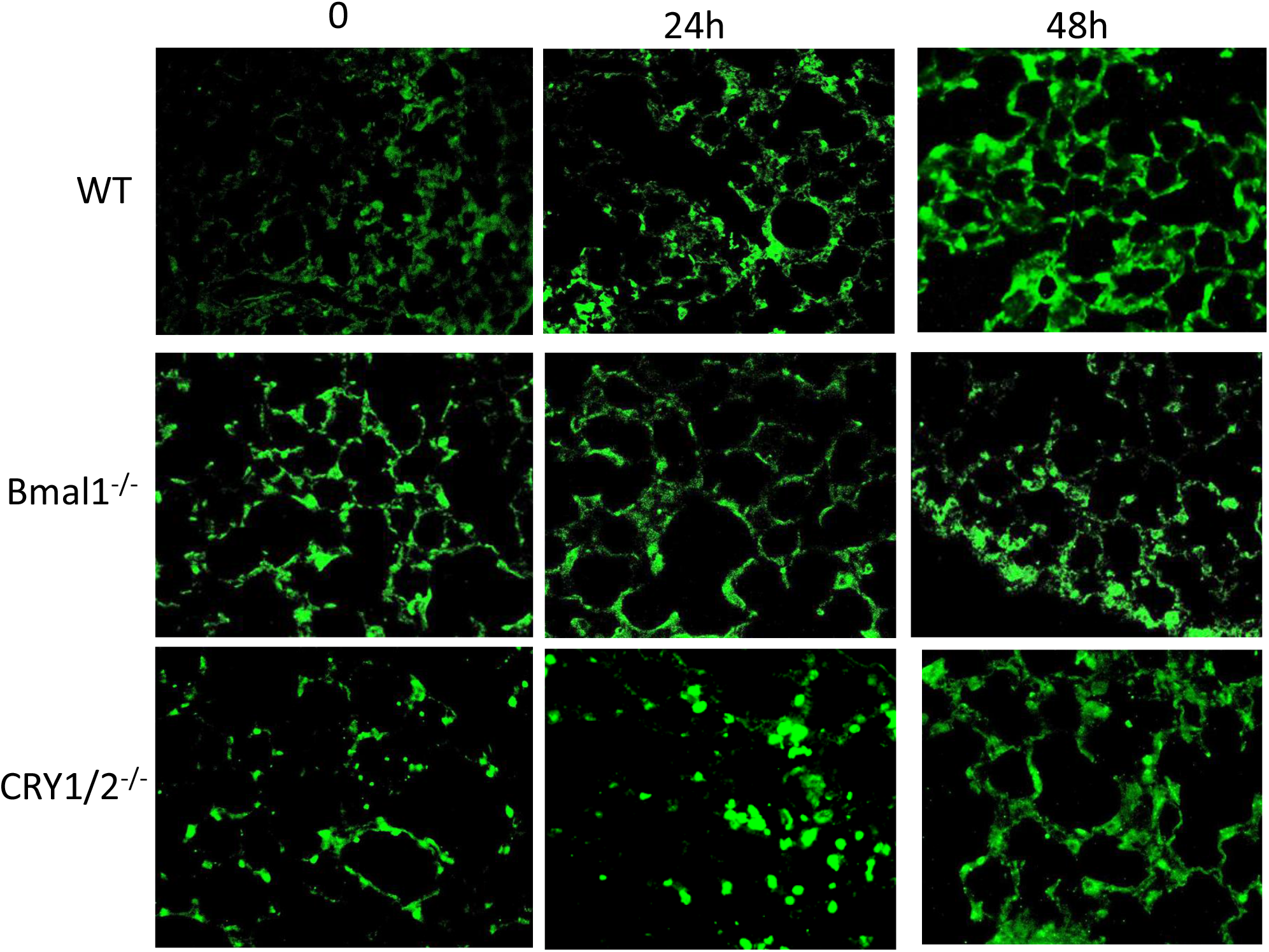
Inflammation in lungs as monitored by immunostaining for Polymorphonuclear Neutrophils (PMN) in lungs of mice (WT, Bmal1 null, CRY1/2 null) after i.t. LPS instillation. Lung sections were immunostained for PMN (Ly6G) by anti-ly-6G6C antibody (also called NIMP). Secondary antibody was tagged to Alexa 488.

**Figure 8 B.**
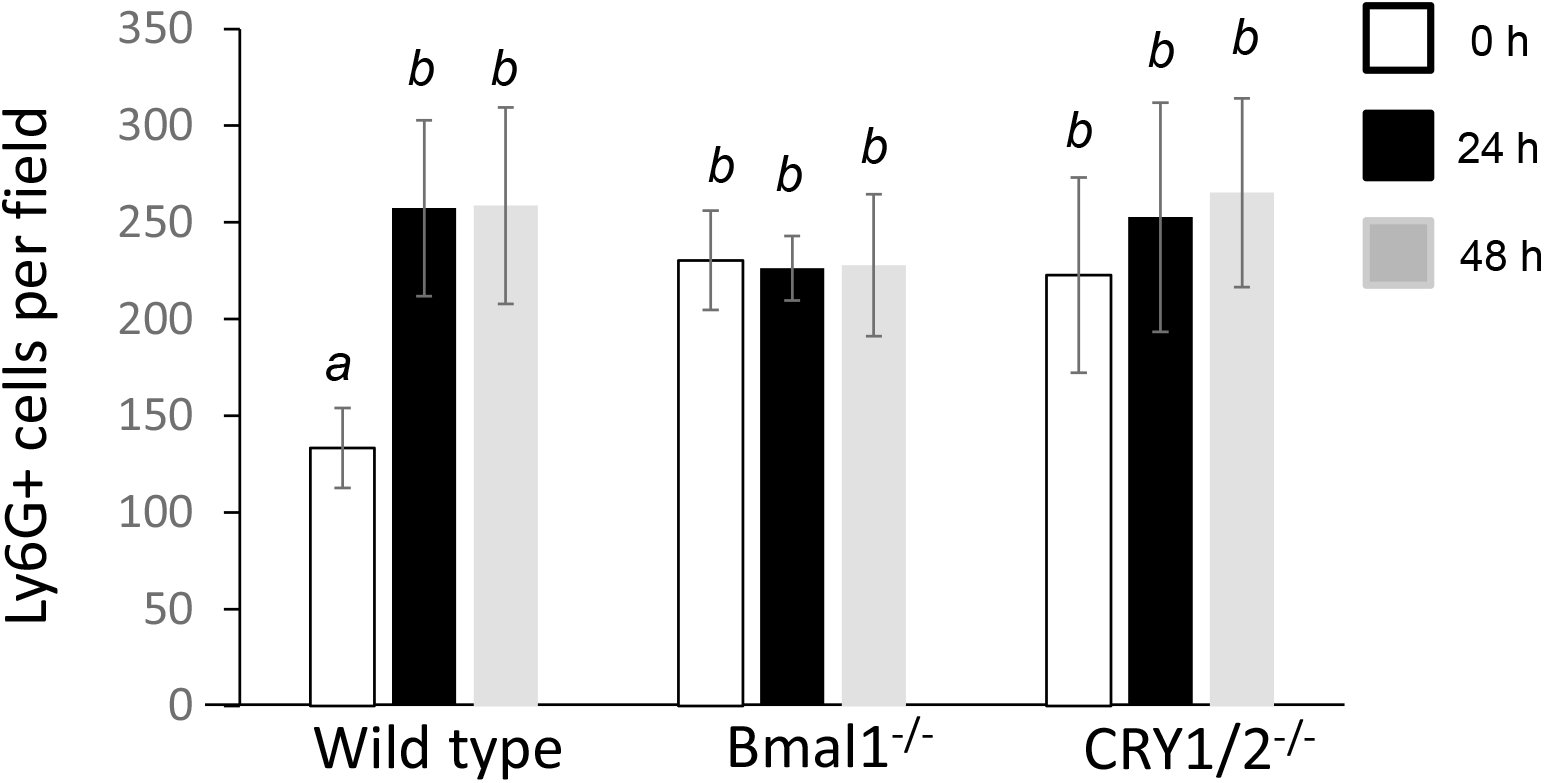
Fluorescence signal (green dots that represent PMN) per field was quantitated over several sections using Metamorph Software. n=3 WT mice and CRY1/2 null and n=2 Bmal null were used. For each mouse, at least 4-5 sections were immunostained and imaged. 3 fields were imaged per section to obtain an average for each section. Images were quantified by semi-automate image analysis that counts particles (green dots). All sections was averaged, and data as expressed as means± SD. Different letters between groups denote statistically significant differences (p<0.05) between those two groups. If two groups share the same letter, then they are not statistically significantly different

**Figure 8 C.**
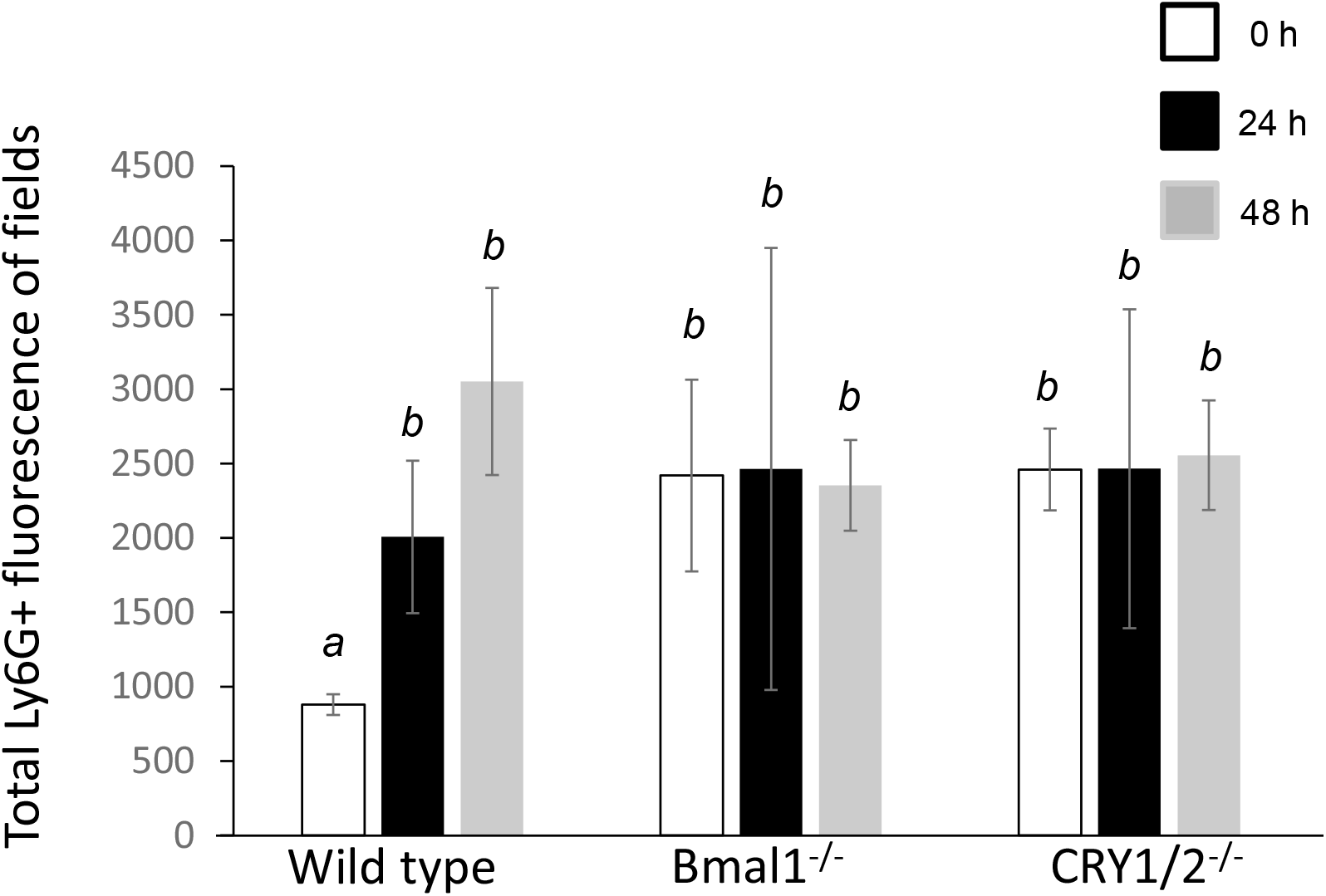
Total PMN activation captured via integrated fluorescence Intensity of Ly6G+. Different letters between groups denote statistically significant differences (p<0.05) between those two groups. If two groups share the same letter, then they are not statistically significantly different

## Discussion

The role of the circadian clock in regulation of inflammation is well known. Conversely, inflammation itself can directly affect circadian rhythmicity [22]. The importance of the clock in inflammation was clear more than half a century ago when studies showed that the mortality rate of mice exposed to the bacterial endotoxin (lipopolysaccharide, LPS) depended on time of exposure [23–26] i.e., LPS challenge given at the end of the rest time resulted in a mortality rate of 80%, while that given in the middle of the active time resulted in a mortality rate of only 20%[23]. Furthermore, bacterial and viral infection has been shown to lead to higher mortality when initiated during the rest period [7, 24, 27]. Similarly, the effect of increased inflammation on dysregulation of the circadian rhythm has been well established; indeed, treatment with IFN-γ reduced the amplitude of the circadian rhythm of cultured rat SCN neurons [28].

To date, the interrelations between inflammation and circadian rhythm have primarily been studied on immune and inflammatory cells [29]. Scant attention has been paid to the endothelium, specifically the pulmonary endothelium, which by virtue of its location serves both as a participant of inflammation signaling (endothelial signaling upregulates moieties that drive adherence and transmigration of immune cells) and as a converging site whereby immune cells (such PMN) adhere to the vessel wall, followed by their transmigration into tissue [8, 9]. Here we monitored the circadian rhythmicity of the pulmonary endothelium with inflammation and evaluated the role of the clock in regulating pulmonary inflammation. As Bmal1 is known to have significant developmental or non-circadian effects, we used both Bmal1 and CRY1/2DKO to investigate the association of pulmonary endothelial inflammation signaling with the clock. Considering the multitude of inflammation driven lung pathologies [21, 30], identifying the interrelation between Bmal1, CRY1/2 and pulmonary inflammation is critical to develop therapeutic strategies for the control of lung inflammation and injury.

A pivotal molecule that drives endothelial inflammation and injury is the NLRP3 inflammasome. It is a multiprotein complex comprised of three basic components: (1) A sensor such as a NOD-like receptor (NLR) (2) the adaptor protein apoptosis-associated speck-like protein containing a caspase-recruitment domain (ASC) and (3) the inflammatory cysteine aspartase caspase 1 [31, 32]. Enhanced NLRP3 expression has been well established as a pivotal event in lung hyper-inflammatory syndromes [21]. NLRP3 signaling is associated with inflammatory and metabolic diseases ranging from arthritis and gout to atherosclerosis and type 2-diabetes [33]. Recently the biological clock was reported to control NLRP3 in certain immune cells and NLRP3 protein amounts were found to be modulated by Rev-erbα activity [34].

Our data show that an inflammatory stimulus [endotoxin (LPS)] resets the circadian rhythm via a NADPH oxidase 2 driven ROS signaling pathway. This suggests that oxidants can affect the phase and amplitude in pulmonary endothelium; other cell types too have shown that the cellular clock is susceptible to ROS [35, 36]. How does ROS exert effects on the circadian rhythm, and how is the pulmonary endothelium affected by this altered amplitude? ROS is well established to upregulate several inflammatory moieties on the pulmonary endothelium; among these is the NLRP3 inflammasome, that we and others have shown to be induced by ROS [17, 37]. We thus reasoned that the alteration of circadian rhythms by ROS could be via NLRP3. We observed that NLRP3 expression which was low in untreated lungs increased after LPS exposure; the expression peaked at 36 h and reduced thereafter. Inflammation (ICAM1 expression and PMN adherence) followed a similar pattern. However, Bmal and CRY null lungs did not show a similar profile of onset, increase and reduction in NLRP3 and inflammation. In pulmonary endothelial cells we observed that inflammation alters the amplitude of the circadian cycle and does not disrupt circadian rhythm in the pulmonary endothelium. Thus, we speculate that the increase in amplitude following LPS may serve an important role in the resolution and subsequent recovery. The fact that we do not see this pattern of initial upregulation and subsequent resolution in mice wherein the clock has been disrupted genetically, argues that with clock disruption, the lung’s ability to resolve and recover from inflammation is impaired. The fact that ICAM-1 and PMN levels were high in lungs of naïve clock mutants and remained high throughout the observed period in contrast to wild type littermates where these levels were resolved to almost control levels, underscore the impact of circadian disruption on inflammation and its resolution.

While the pro-inflammatory phenotype of circadian disruption has been described and studied extensively, we report here the features unique to this regulation in the pulmonary endothelium. To the best of our knowledge this is the first of this kind of study to demonstrate a role for altered circadian clock in resolving pulmonary endothelium.

Taken together, we show that NOX2 driven ROS production resets circadian rhythms in the pulmonary endothelial cells of WT mice; this process is mediated via NLRP3. As expected, increased NLRP3 facilitates lung inflammation. Circadian genes regulate inflammation post LPS treatment via resolution of NLRP3; lack of Bmal1 and CRY1/2 blocks this regulation and prevents the resolution of inflammation (**Schema, Figure 9**).

**Figure 9.**
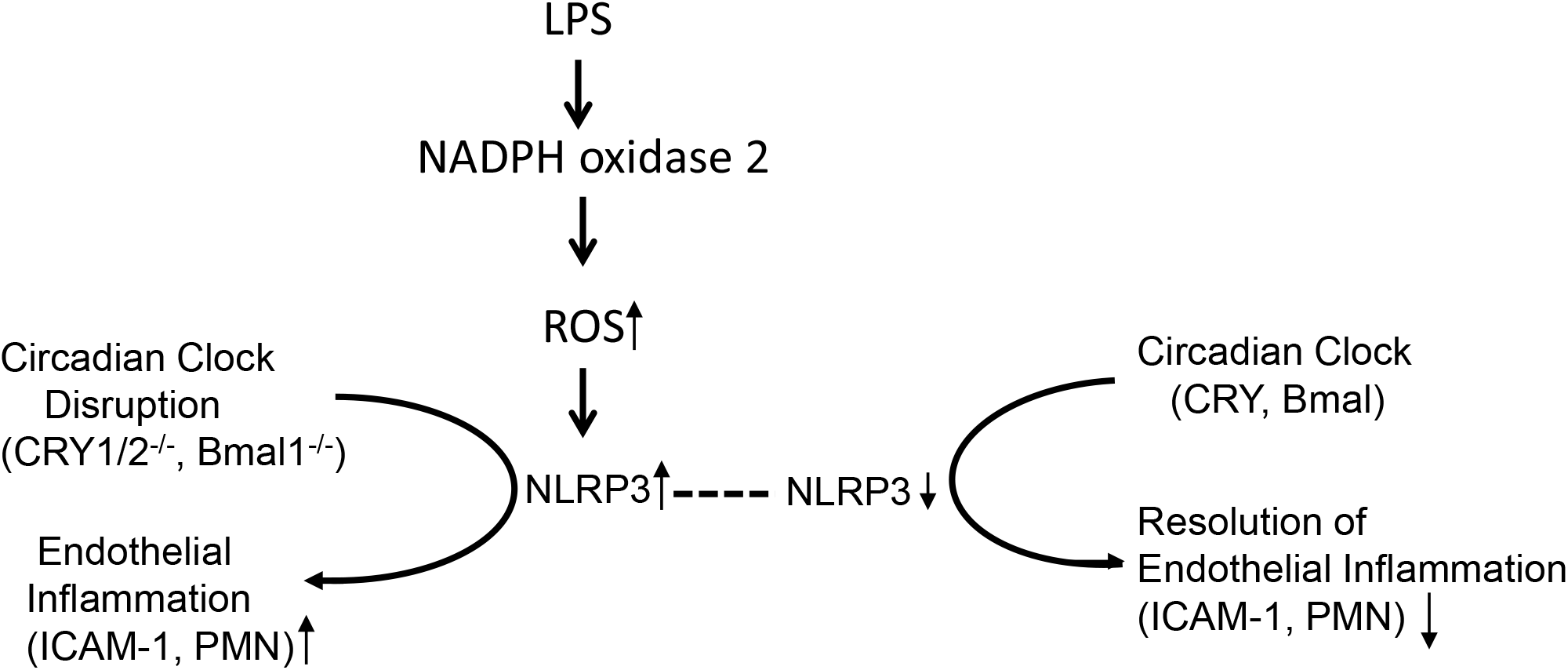
Schema showing the association of NOX2-ROS, NLRP3 and circadian genes with pulmonary endothelial inflammation. Circadian rhythm regulates inflammation via NLRP3. In the absence of circadian genes, NLRP3 and inflammation are dysregulated.

## Acknowledgements

This study was in part supported by NIH R56 HL139559 awarded to SC. AS is a HMMI investigator.

